# Design and Performance Evaluation of a Wearable Passive Cable-driven Shoulder Exoskeleton

**DOI:** 10.1101/2020.05.29.096453

**Authors:** Morteza Asgari, Elizabeth A. Phillips, Britt M. Dalton, Jennifer L. Rudl, Dustin L. Crouch

**Affiliations:** Department of Mechanical, Aerospace, and Biomedical Engineering, University of Tennessee, Knoxville, TN 37996 USA; Brain and Spine Institute, Department of Rehabilitation Services at the University of Tennessee Medical Center, Knoxville, TN 37920 USA; Department of Mechanical, Aerospace, and Biomedical Engineering University of Tennessee, Knoxville, TN 37996 USA

**Keywords:** Assistive technology, Exoskeletons, Human-robot interaction, Shoulder disability

## Abstract

The mechanical assistance provided by exoskeletons could potentially replace, assist, or rehabilitate upper extremity function in patients with mild to moderate shoulder disability to perform activities of daily living. While many exoskeletons are “active” (e.g. motorized), mechanically passive exoskeletons may be a more practical and affordable solution to meet a growing clinical need for continuous, home-based movement assistance. In the current study, we designed, fabricated, and evaluated the performance of a wearable, passive, cable-driven shoulder exoskeleton (WPCSE) prototype. An innovative feature of the WPCSE is a modular spring-cam-wheel system that can be custom designed to compensate for any proportion of the shoulder elevation moment due to gravity over a large range of shoulder motion. The force produced by the spring-cam-wheel system is transmitted over the superior aspect of the shoulder to an arm cuff through a Bowden cable. The results from mechanical evaluation revealed that the modular spring-cam-wheel system could successfully produce an assistive positive shoulder elevation moment that matched the desired, theoretical moment. However, when measured from the physical WPCSE prototype, the moment was lower (up to 30%) during positive shoulder elevation and higher (up to 120%) during negative shoulder elevation due primarily to friction. Even so, our biomechanical evaluation showed that the WPCSE prototype reduced the root mean square (up to 35%) and peak (up to 33%) muscular activity, as measured by electromyography, of several muscles crossing the shoulder during shoulder elevation and horizontal adduction/abduction movements. These preliminary results suggest that our WPCSE may be suitable for providing movement assistance to people with shoulder disability.

## I. Introduction

Shoulder disability is a global health burden [1, 2] associated with several common orthopedic and neurological disorders, such as rotator cuff tear [3], peripheral nerve injury [4], muscle atrophy [5], and stroke [6]. Many people with shoulder disability have mild to moderate disability, meaning that they can still move their shoulder but with lower strength and range of motion than that of a healthy shoulder. Furthermore, people with shoulder disability often have trouble elevating the shoulder or holding the arm up against gravity. Such functional deficits can make it hard for individuals to perform various activities of daily living, particularly those activities whose net shoulder joint moments are dominated by gravity [7]. Consequently, shoulder disability can negatively impact an individual’s quality of life, selfesteem, and independence.

For people with shoulder disability, there is a clinical need for medical devices that can provide continuous, home-based movement assistance to compensate for gravity at the shoulder. Gravity compensation can assist upper extremity motor function by reducing the joint moments that muscles need to generate for a given movement [8]. We previously showed that anti-gravity assistance reduces activations of muscles that mainly contribute to positive shoulder elevation [9]. By reducing muscle activations, gravity compensation at the shoulder could help people perform upper extremity tasks with less effort and reduce muscle tissue loads. Additionally, for people with shoulder disability, gravity compensation could enhance shoulder strength and range of motion.

A wide variety of assistive and rehabilitative exoskeletons (sometimes called orthoses) have been developed for people with disability. Exoskeletons, which are typically worn on and transmit forces to the body [10], generate forces either actively or passively. Active exoskeletons, such as ARMin III [11], CADEN-7 [10], and CAREX [12], use motors or other powered actuators to generate the forces that are applied to the user. Active exoskeletons can provide more flexible assistance since the forces they generate can be modulated by control software. However, they are relatively heavy, large, expensive, and complex because they require motors, power sources (e.g. batteries), and computer hardware. These features make active exoskeletons less wearable and portable, limiting their application to clinical and laboratory settings.

Passive exoskeletons generate assistive forces using counterweights, rubber bands, or springs. The assistive forces generated by passive exoskeletons are fixed according to their mechanical design and cannot be modulated in real time, as with active exoskeletons. However, passive exoskeletons are appealing for providing continuous, home-based movement assistance since they do not require electromechanical hardware (e.g. motors, batteries). Thus, passive exoskeletons are potentially more lightweight, lower cost, easier to maintain, wearable, and portable than active exoskeletons.

Existing *wearable* passive shoulder exoskeletons compensate for gravity by releasing and storing energy from elastic springs as the shoulder is elevated and lowered, respectively [13]. Commercially available wearable passive shoulder exoskeletons, intended for occupational or industrial settings, include: EksoVest (Exo Bionics Holdings, Inc.), SuitX (US Bionics, Inc.), and AirFrame (Levitate technologies, Inc.). However, these existing exoskeletons may not be suitable to assist activities of daily living for people with shoulder disability because they are designed for application-specific and high-load tasks (e.g. overhead drilling and heavy lifting) [14, 15]. They also typically have a high profile (i.e. they extend out far from the user’s body) because they incorporate rigid linkages and mechanical joints.

Some passive devices have been developed for clinical applications, but most are not wearable. Early devices, such as passive mobile arm supports that mount on a wheelchair or wall [16, 17], compensate for gravity using counterweights [18] or a parallelogram with a spring-cam mechanism [19]. The Wilmington Robotic Exoskeleton (WREX), a more recent passive mobile arm support [20], uses elastic bands to compensate for gravity at the shoulder and elbow joints. The WREX can be mounted on a body-worn back brace or a wheelchair. These passive devices are limited to provide support only at certain shoulder angles and are unproven to provide modulated gravity compensation over the shoulder’s entire range of motion.

The goal of this study was to design, fabricate, and preliminarily evaluate the performance of a new wearable, passive, cable-driven shoulder exoskeleton (WPCSE) whose features (described in Section II.A) make it suitable for providing continuous, at-home assistance to patients with mild to moderate shoulder disability. We developed a physical benchtop model that allowed us to quantitatively evaluate the mechanical performance of the WPCSE. Additionally, we performed an experiment with four able-bodied participants to evaluate effect of the WPCSE on shoulder muscle activity during repetitive shoulder elevation and horizontal adduction/abduction movements. We hypothesized that activations of muscles crossing the shoulder would be the same or lower with the WPCSE than without.

## II. Design of the WPCSE

### A. System Concept

Our WPCSE compensates for gravity at the shoulder using an elastic spring whose force is transmitted to the arm through a Bowden cable that passes over the superior aspect of the shoulder (Fig. 1). The spring is stretched when the arm is beside the body and pulls on the Bowden cable as the arm elevates.

**Fig. 1.**
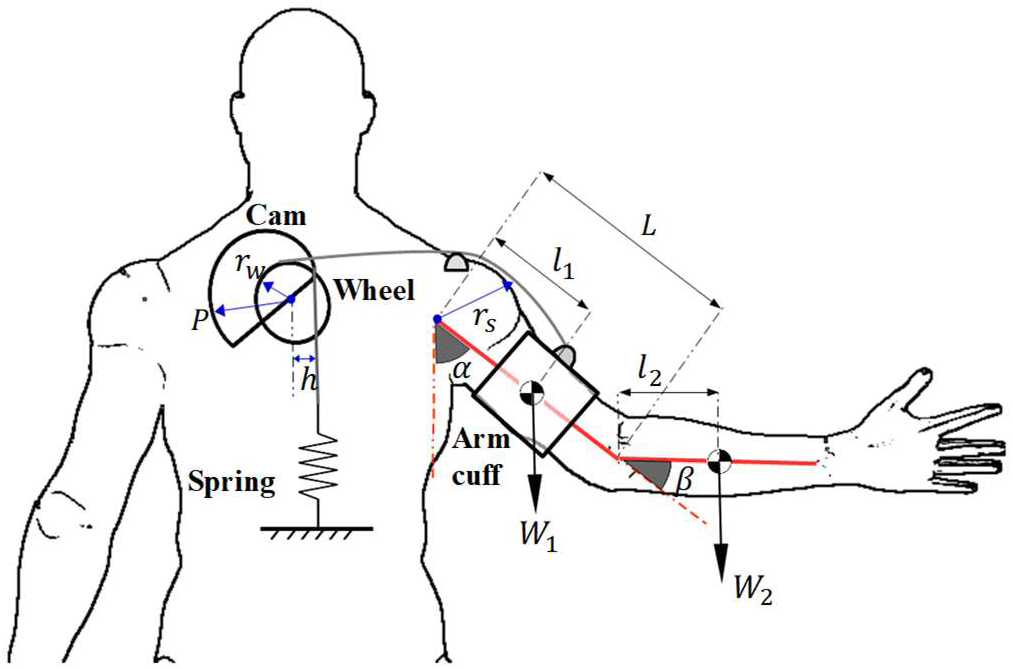
Schematic diagram of shoulder exoskeleton system concept (posterior view on right arm) can maintain a low profile.

As described in section II.B. below, as the shoulder elevation increases from 0° to 90°, the magnitude of the negative shoulder elevation moment due to gravity increases. To provide increasing assistance in proportion to the increasing gravity moment, we incorporated a cam-wheel component between the spring and arm that functions as a gearing mechanism to modulate the cable pulling force. The cam-wheel component consists of a variable-radius cam and a constant-radius wheel that are fixed to each other and rotate together about the same axis. The variable-radius cam attaches to the spring via a wrapping rope, while the constant radius wheel connects to the arm via the Bowden cable. All WPCSE components are mounted on one of two semi-rigid body fittings, a back brace and an arm cuff.

The proposed exoskeleton improves on previous wearable passive shoulder exoskeleton designs in several ways. First, it provides mechanical assistance through a soft, cable-driven mechanism instead of rigid links, which permits us to (1) place force-generating components (i.e. the cam-wheel and spring) away from the shoulder joint, (2) assist shoulder elevation without limiting shoulder axial rotation and elevation plane movements, and (3) reduce the exoskeleton’s weight. Second, the cam-wheel gearing component enables us to tune the gravity-compensating force over a wide range of shoulder motion, making the exoskeleton more suitable for supporting a wide range of static postures and dynamic movements. With our compact spring-cam-wheel system design, the exoskeleton can maintain a low profile.

### B. Estimation of Shoulder Elevation Moment due to Gravity

The magnitude of the gravity moment about the shoulder depends on the upper extremity posture, while the orientation of the gravity moment depends on the orientation of the torso with respect to gravity. In our design calculations, we assumed that the torso is upright and aligned with the gravity vector. In this case, gravity will primarily generate a moment about the shoulder in the direction of negative shoulder elevation. The shoulder gravitational moment caused by the weight of the arm and forearm/hand segments is:

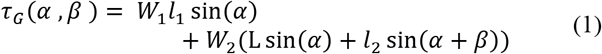

Where *W*_1_ is the weight of the arm; *W*_2_ is the sum of the forearm and hand weight; *l*_1_ is the approximate distance from the glenohumeral joint center to the arm’s center of gravity; *l*_2_ is the approximate distance from the elbow joint center to the center of gravity of the combined forearm-hand segment; L is the length of the arm segment; and *α* and *β* are the shoulder elevation and elbow flexion angles, respectively (Fig. 1). Equation (1) shows that the shoulder elevation moment is coupled with the gravitational moment at the elbow joint and increases nonlinearly as the shoulder elevates from 0° to 90°. We estimated the value of *τ_G_* in (1) based on anthropometric data of a 50^th^ percentile male (age 27.45(5.64) *years*) and assuming a fully extended elbow [21, 22].

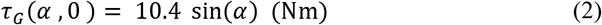

### C. Selection of Elastic Spring Component

The force-generating components of passive devices are typically either zero-free-length springs, constant-force springs [23], or rubber bands [24]. For the shoulder joint, which has the largest range of motion of all human extremity joints, an ideal passive element must be able to generate force over a large range of deflection. This eliminates extension springs as design options because they can only stretch up to a maximum length equal to their free length [19]. Another type of passive element is a constant-force spring, which basically includes a prestressed coil of a flat, elastic strip that produces a nearly constant force when its end is translated over a long distance. Constant-force springs, however, require space to be unwound enough to develop their full force.

For our WPCSE, we used another elastic component called a constant-torque spring (Fig. 2). Compared to constant-force springs, constant-torque springs are more compact since their deflection is achieved by rotating a set of spools. In a constant-torque spring, a coiled, elastic metal strip is wrapped over a storage spool. The metal strip is stretched by wrapping and rotating it around an output spool in the opposite direction in which it is coiled. Energy is released from the spring when the metal strip unwinds from the output spool back onto the storage spool. When the spring is stretched, it generates constant torque *τ_spring_* about the axis of the output spool [25]. We converted the torque into a tension force in a wrapping rope by attaching a pulley to the output spool (Fig. 2).

**Fig. 2.**
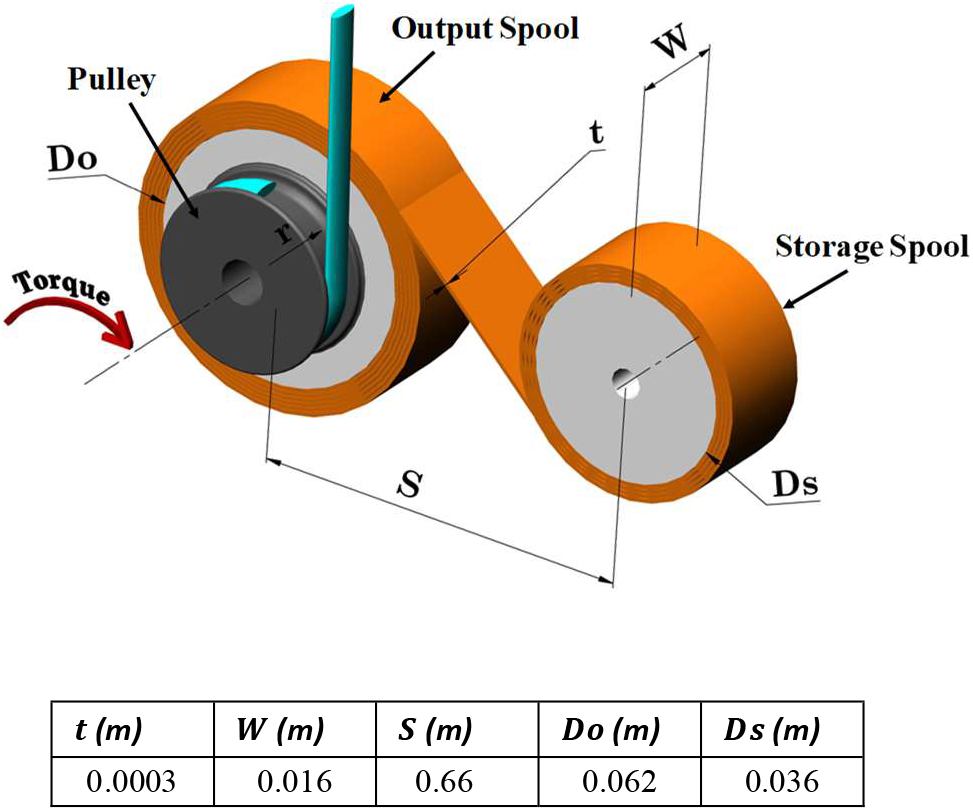
Schematic view of a constant-torque spring. The geometrical properties of the selected spring assembly are listed in the table.

Based on the gravity moment in (2) and initial design work, we selected a commercially available constant-torque spring that can nominally produce a torque, *τ_spring_* = 0.84 + 0.08 *Nm* when is wound 10 turns around the output spool, according to the manufacturer (SV12J192, Vulcan Spring & Manufacturing, Telford, PA, USA). The spring material is 301 Stainless steel and the design parameters recommended by the manufacturer are shown in the table in Fig. 2.

The torque generated by a constant-torque spring is higher during the stretching phase (i.e. spring winding onto the output spool) than during the recoil phase (i.e. spring winding back onto the storage spool). We opted to perform the design of WPCSE based on the torque generated during the recoiling phase during which it assists positive shoulder elevation movements. The torque, *τ_spring_*, that we measured (see section III.A.) from the constant-torque spring during the recoiling phase was 0.67 ± 0.02 *Nm*.

### D. Development of Cam-wheel Profile

The cam-wheel is the key component of the WPCSE that adjusts the cable pulling force to output the desired increasing positive shoulder elevation moment with increasing shoulder elevation angle. (Fig. 3). The rotation of the wheel (*φ*) is related to the shoulder elevation angle (*α*), and can be described as (3):

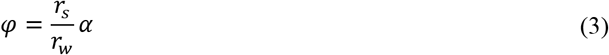

*r_w_* represents the radius of the constant-radius wheel and *r_s_* is the moment arm of the cable around the glenohumeral joint. Theoretically, the WPCSE can be designed to generate an assistive positive shoulder elevation moment *τ_E_* that counteracts some proportion, *k*, of the gravity moment, *τ_G_*(*α*, 0). Here, we set *k* to a constant value of *k* = 1/4. This value was based on our preliminary study in which, for *k* = 1/2, some users needed to actively lower their arm with the WPCSE [26].

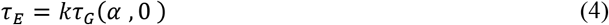

**Fig. 3.**
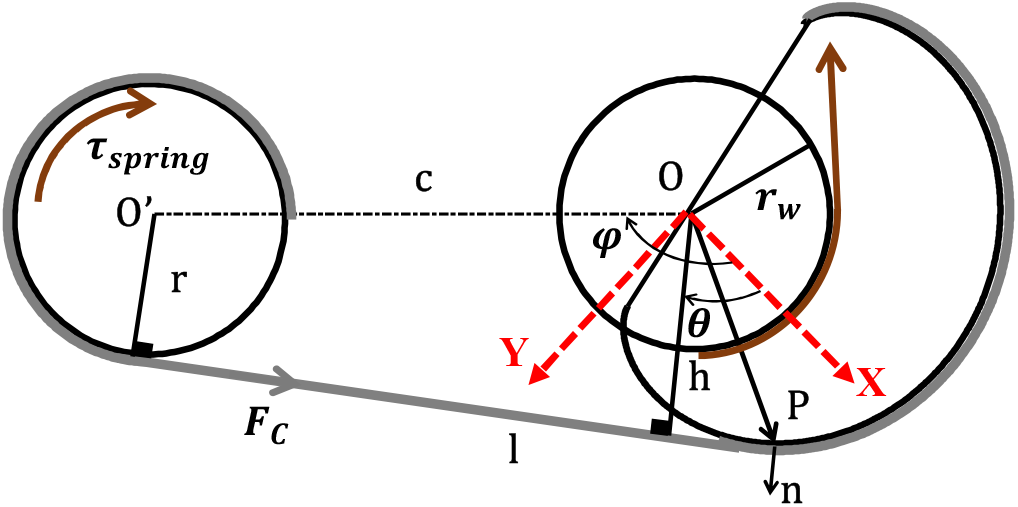
Schematic representation of the spring-cam-wheel system.

For the cam-wheel to be statically balanced, assuming no loss due to friction:

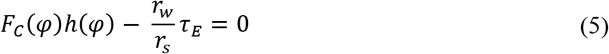

Where *F_C_*(*φ*) is the tension force in the wrapping rope that connects the constant-torque spring to the variable-radius cam, and *h*(*φ*) is the moment arm of the wrapping rope with respect to the cam’s axis of rotation.

We first determined the form of *h*(*φ*) in (5) to calculate the final geometry of the cam. The value of *r_s_* was set to 0.06*m*, which we estimated from a computational musculoskeletal model [27]. The cable tension force was calculated by dividing *τ_spring_* by the radius *r* of the pulley which is attached to the output spool. To determine the form of *h*(*φ*) and values of *r_w_* and *r*, we performed a constrained, global numerical optimization using the *GlobalSearch* function in MATLAB (MathWorks, Inc., Natick, MA). The optimization identified parameter values that minimized the sum of the squared error between the actual (*τ_E_*) and desired 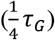 assistive shoulder elevation moment by the exoskeleton, 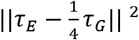. From among five forms for *h*(*φ*), including linear, quadrature, polynomial, power, and sinusoidal functions, that we tested in our global optimization procedure, we chose a sinusoidal form because it generated the lowest error [26]. This makes sense since the form of *τ_G_* in (2) is also sinusoidal.

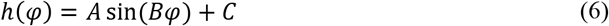

The design parameters computed during the global optimization procedure are displayed in Table I.

**TABLE I.** Optimized Design parametrs

| $r_w$ (m) | $r$ (m) | $A$ | $B$ | $C$ |
| --- | --- | --- | --- | --- |
| 0.030 | 0.030 | -0.057 | -0.492 | 0.000 |

We then determined the geometrical profile of the cam analytically using a complex form of the loop closure method [28]. The coordinate of the contact point, *P*, between the wrapping rope and the cam surface (Fig. 3), can be represented as:

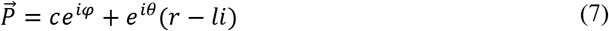

Where *c* is the center-to-center distance between cam-wheel and the spring output spool; *l* is the length of the unwrapped portion of the wrapping rope; and *θ* is the angle of the moment arm. The angle *φ* can be written as a function of the moment arm *h*(*φ*):

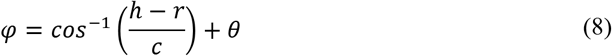

We set *c* = 0.010*m* to have the final integrated spring-cam-wheel system as compact as possible while still maintaining clearance between the cam-wheel component and the constant-torque spring. The value of *l* was derived using the contact condition between the cable and the cam 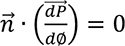. This leads to the following equation to calculate *l*.

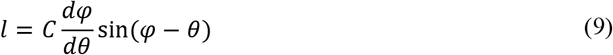

Where 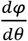 can be determined by taking the derivative of (8):

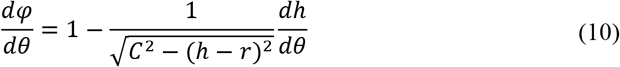

Equations 7 to 10 are sufficient to develop the final profile of the cam-wheel (Fig. 4A). The coordinates of three hundred points representing the cam profile were obtained to develop its computer-aided design (CAD) model (Fig. 4B).

**Fig. 4.**
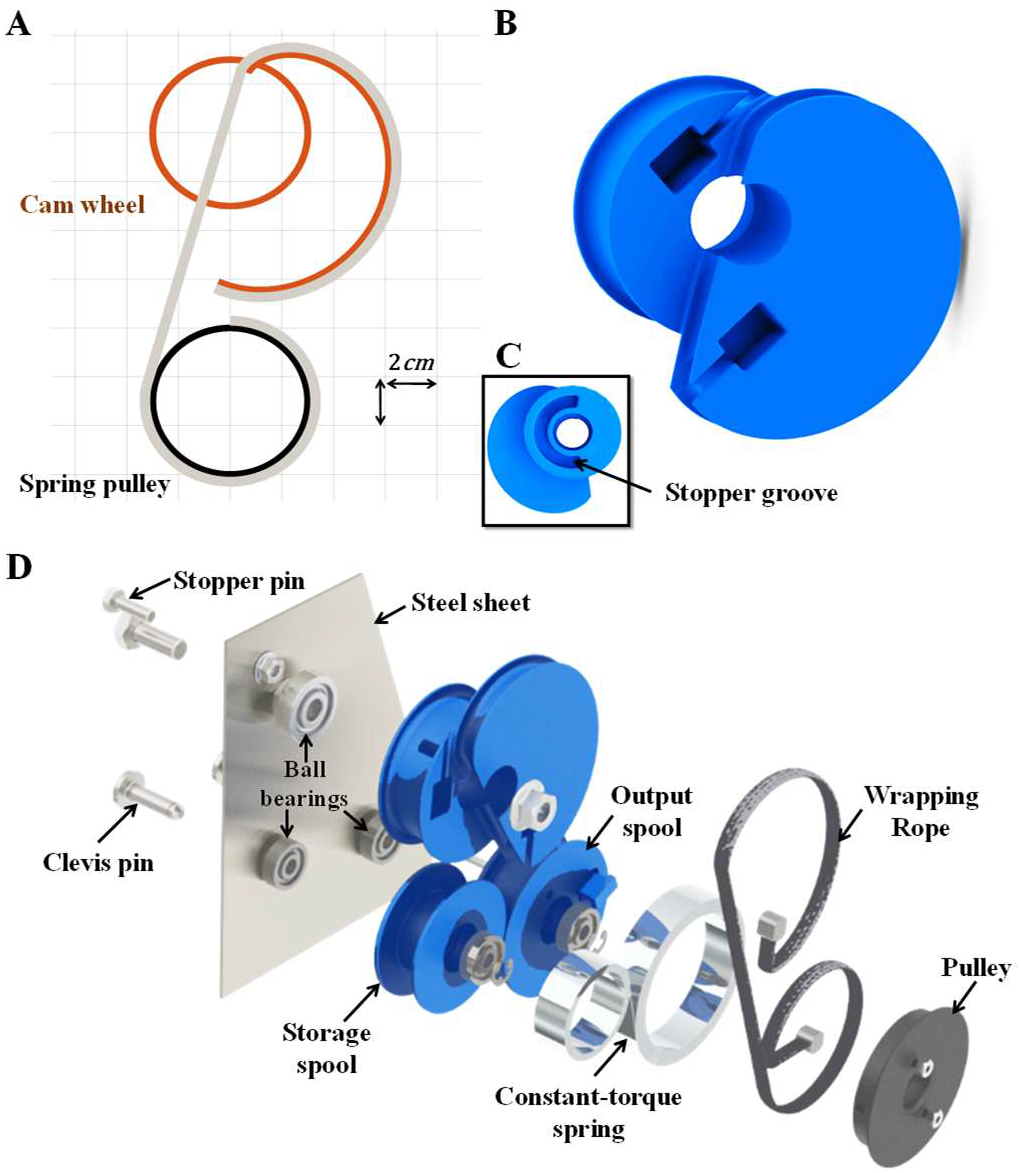
A) Profile of the cam-wheel. B) The CAD model of the cam-wheel. C) Posterior view of the cam-wheel. D) Exploded view of the spring-cam-wheel assembly.

### E. Exoskeleton Prototype

Based on the design parameters and dimensions defined in previous sections, we fabricated a prototype of the WPCSE using off-the-shelf and 3D-printed components, as described below.

#### 1) Spring-cam-wheel Assembly

The CAD models of the cam-wheel, spring spools, and pulley were developed in Solidworks software (2018, Dassault Systémes SolidWorks Corp., MA, USA). From the CAD models, the parts were 3D printed in ABS plastic (Fig. 4B, 4C, 4D). The constant-torque spring and output pulley were assembled and installed on a 304-alloy stainless-steel sheet (14*cm*×17*cm*×0.16*cm*) using Clevis pins and retainer rings. To facilitate the rotation of the spools, two deep-groove ball bearings (688-ZZ 8*mm*×22*mm*×7*mm*) were pressed in each spool. The cam-wheel was also equipped with a single row ball bearing (Koyo, EE4C3) at its center, and bolted to the stainless sheet. A non-stretching, solid braid nylon rope (9.50 *mm* thickness) attached the cam to the spring pulley. To prevent the constant-torque spring from suddenly winding onto the storage spool, a mechanical stopper was incorporated in the output spool to limit its rotating to 350°. The stopper simply consisted of a pin that was bolted to the stainless-steel sheet and fit inside a circular groove on the underside of the output spool. A similar stopper was placed beneath the cam-wheel to limit its rotation such that the WPCSE could apply force to the arm for shoulder elevation angles between 0° and 90°. Figure 4D shows an exploded view of the power source assembly.

#### 2) Fittings

The WPCSE prototype consisted of two body fittings, a back brace and an arm cuff, on which other components were mounted. We custom-made a thermoplastic back brace (acrylic sheet) by casting a person’s torso with plaster gauze and fabricating positive and negative molds. The back brace was attached to a back-support vest that straps across the shoulder and around the waist (Fig. 5A). The inside of the back brace was covered with padding to improve fit and comfort. A thermoplastic cuff was attached to an adjustable, slide-on elbow support that straps to the arm both above and below the elbow joint (Fig. 5A). This prevented the arm cuff from sliding proximally while still permitting elbow flexion.

**Fig 5.**
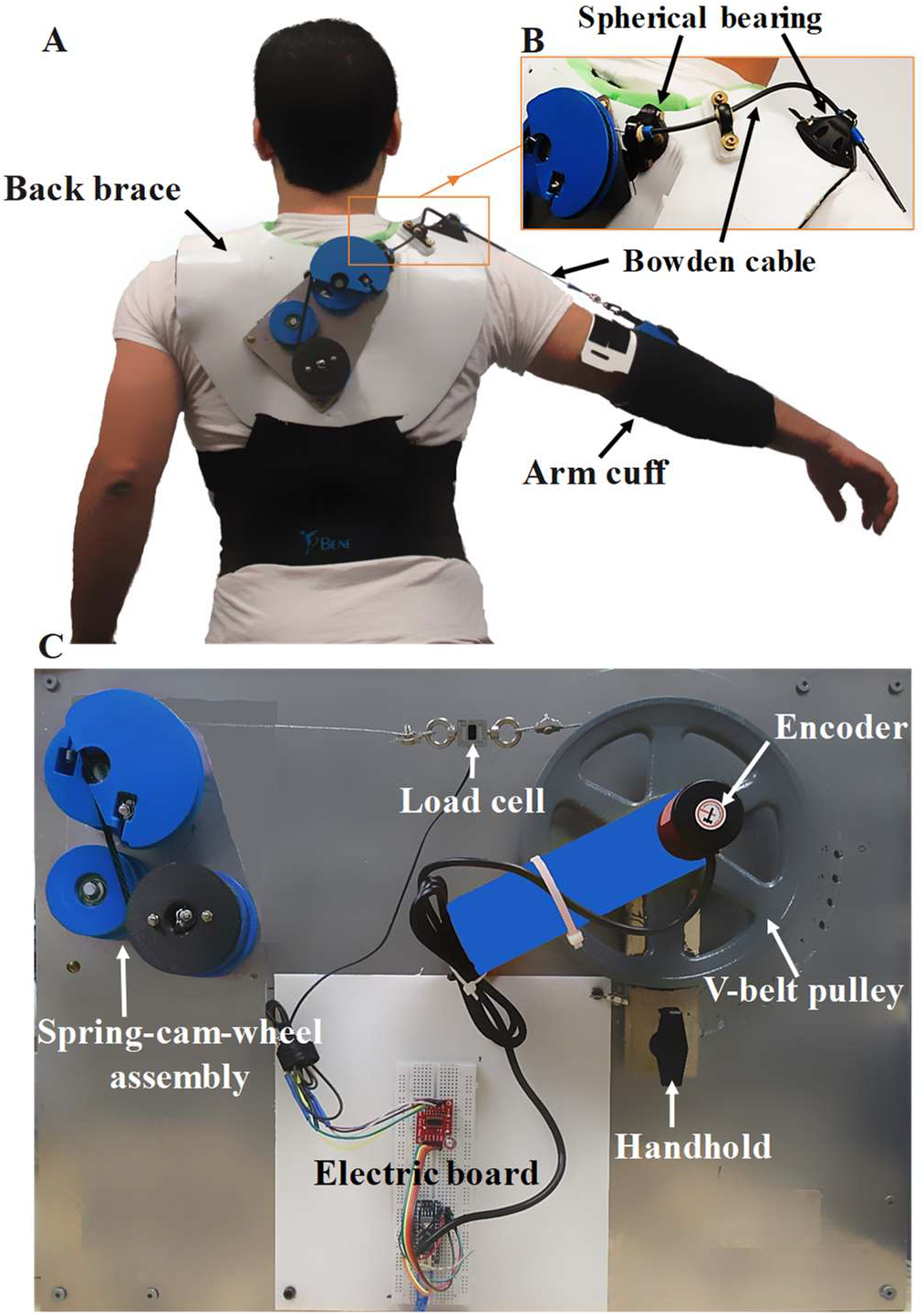
A) WPCSE prototype. B). Bowden cable routing path over the shoulder C) Benchtop prototype.

#### 3) Cable Routing

A Bowden cable was used to transmit the mechanical force from the spring-cam-wheel assembly to the arm cuff. A Bowden cable consists of an inner cable (commonly of steel) and an outer casing (a composite of helical steel wire and plastic outer sheath). The inner wire wrapped around the wheel on one end and connected to the arm cuff on the other end using a spring snap.

The Bowden cable casing was clamped at each end and in the middle by self-aligning spherical bearings (maximum pivot angle = 60°) that were bolted to the back brace (Fig. 5B). Clamping the casing was required so that so that the inner wire could move relative to it and transmit the pulling force. The spherical bearings permitted the cable to pivot as its orientation changed with shoulder rotation. The path of the Bowden cable was designed to minimize friction (i.e. have as few bends with as large radii as possible). A small (~5cm) length of casing extended distally from the bearing mounted over the shoulder to enable the inner cable to curve smoothly over the shoulder.

#### 4) Prototype Specifications

The final prototype of the WPCSE weighed 1.82 *kg*, of which 0.83 *kg* was due to the spring-cam-wheel assembly, 0.82 *kg* was due to the back brace, and 0.17 *kg* was due to the arm cuff. The spring-cam-wheel system had the major weight of the exoskeleton mostly due to its metallic components. The overall weight of the WPCSE could be reduced further by selecting a spring with an optimal (i.e. shorter) strip length and trimming the shape of the steel board and thermoplastic brace.

The WPCSE prototype was designed so that it could be donned/doffed independently by the user. However, the length of the inner wire of the Bowden cable needed to be adjusted once for each user according to their anthropometry. This can be done by either adjusting the length of the strap attached to the arm cuff or changing the length of the inner wire itself.

### F. Benchtop Setup

In addition to the WPCSE prototype, we developed a benchtop prototype (Fig. 5C) to perform mechanical validation of our theoretical model. The benchtop prototype consisted of a v-belt pulley (radius = 0.08*m*) representing the Bowden cable wrapping over the shoulder; the pulley had an embedded handhold so that it could be rotated manually to mimic shoulder elevation movements. Depending on the test (Section III.A), the v-belt pulley was connected to either the spring-cam-wheel assembly or to the WPCSE. A rotary encoder (LPD-3806-600bm-G5-24c, GTEACH, China) was installed on top of the v-belt pulley to measure its rotation angle for computing the mimicked shoulder elevation angle. An s-type load cell (ATOLC-S04, 50kg capacity, ATO, Diamond Bar, CA, USA) was also placed along the cable path to measure its pulling force. An Arduino Nano board was programmed to compute the shoulder elevation moment and angle from the measured force and angle, respectively.

## III. EXPERIMENTS

### A. Mechanical Performance Evaluation

To validate our theoretical model of the shoulder elevation moment generated by the exoskeleton (i.e. Equation (4)), we measured the torque generated by both the spring-cam-wheel assembly alone and the entire WPCSE using the benchtop prototype. The spring-cam-wheel assembly and WPCSE were essentially the same except that the WPCSE also included the Bowden cable and its routing path on the back brace. The WPCSE was placed on the fabricated positive mold of the torso and fixed to a workbench. The side of the Bowden cable that attaches to the arm cuff was connected to the v-belt pulley. For the spring-cam-wheel assembly, we fixed it on the benchtop board and connected the wheel to the v-belt pulley (Fig. 5C). The v-belt pulley was manually rotated from 0° to 67.5° to simulate the shoulder elevation from 0° to 90°. The angle difference was due to the difference between the v-belt pulley radius and the estimated moment arm of the Bowden cable about the shoulder.

The v-belt pulley was rotated manually counterclockwise (i.e. simulated positive shoulder elevation) and clockwise (i.e. simulated negative shoulder elevation), three times at a slow speed (i.e. quasi-static condition). Cable pulling force and pulley rotational angle data were collected at 20 Hz. The shoulder elevation moment was calculated by multiplying the cable pulling force by the radius of the v-belt pulley. Measured 4 pulley rotation angles were multiplied by 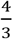 (pulley-to-shoulder moment arm ratio) to compute the simulated shoulder elevation angles. The final moment-angle relationships for the spring-cam-wheel and WPCSE were obtained by resampling the simulated positive and negative shoulder elevation phases into 50 points each and averaging across the three repetitions.

With a similar approach, we quantified the torque-angle relationship for the constant-torque spring alone. The spring was installed on the benchtop board, and the output pulley was attached to the v-belt pulley after winding the spring strip 10 turns around the output spool. The torque of the spring was calculated by multiplying the cable tension force by the radius of output pulley (0.03*m*). We computed the spring’s torque-angle relationship as described above for the spring-cam-wheel and WPCSE.

### B. Biomechanical Performance Evaluation

#### 1) Participants

We evaluated the effect of the WPCSE on the neuromuscular activity of muscles crossing the shoulder in four able-bodied participants (3 males and 1 female, age = 27.25(4.45) *years*, height = 1.74(0.07) *m*, weight = 68.40(5.12) *kg*). Eligible participants were 18 to 70 years old with no self-reported history of upper limb pain and injury within the previous year.

#### 2) Instrumentation and Experimental Procedure

The experimental procedure was approved by the institutional review board at the University of Tennessee, and all participants provided their informed consent. Participants were fitted with the WPCSE and asked to perform shoulder movements with and without the WPCSE (Fig. 6A). The movements were shoulder elevation in the sagittal and frontal planes (Fig. 6B, 6C,), and shoulder adduction/abduction in the horizontal plane (Fig. 6D). For each trial, participants moved the shoulder from 0° to 90° and back to 0° in the relevant plane (Fig. 6). The trials were performed with the elbow fully extended, at the participant’s preferred speed, and with a brief (1-2 seconds) pause at the upper (90°) and lower (0°) limit of movement. In each trial, participants performed 10 continuous repetitions of each shoulder movement while sitting upright on a chair. We randomized the order of exoskeleton condition (with and without WPCSE) and, within each exoskeleton condition, randomized the order of shoulder movements. Participants rested between trials for about 30 seconds to reduce the likelihood of fatigue.

**Fig. 6.**
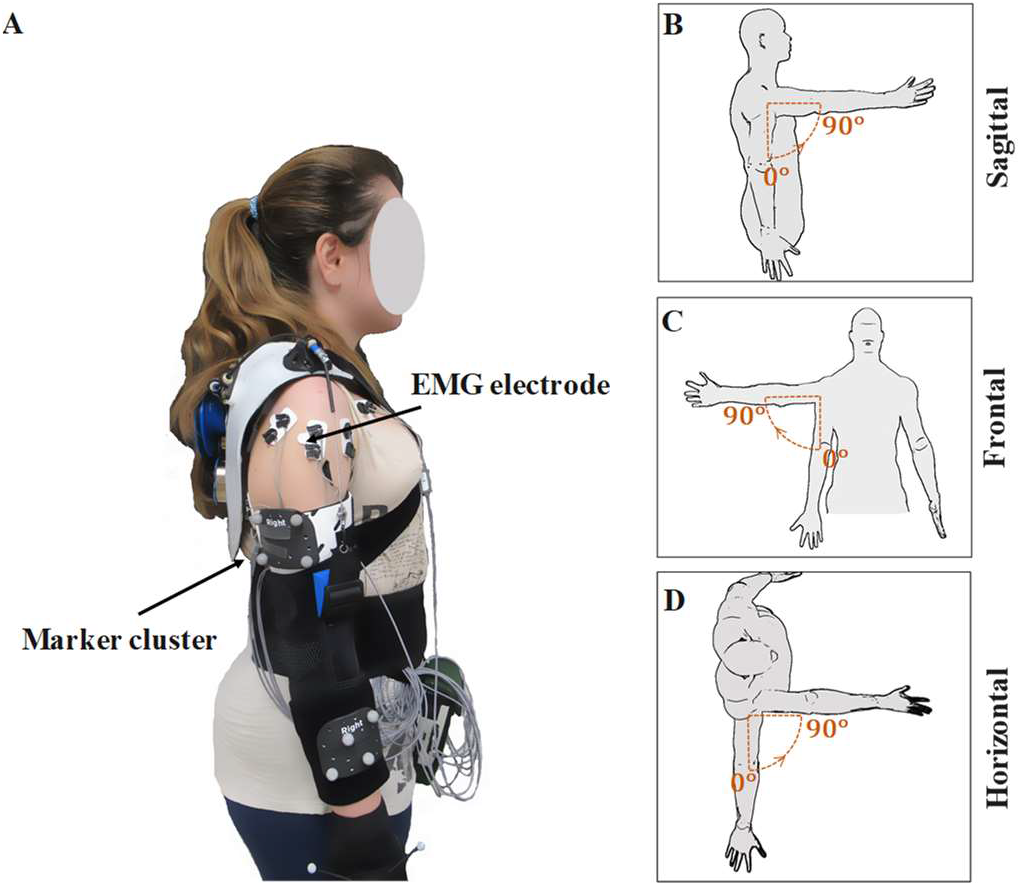
A) A participant with WPCSE, EMG electrodes, and reflective marker clusters. B). Shoulder elevation in the sagittal plane. C) Shoulder elevation in the frontal plane. D) Shoulder adduction/abduction in the horizontal plane.

During the trials, we synchronously measured electromyograms (EMG) and kinematics of the participants. We recorded EMG from several muscles crossing the shoulder, including anterior (AD), middle (MD), and posterior deltoid (PD), pectoralis major (PM), latissimus dorsi (LD), infraspinatus (ISP), trapezius (TRAP), biceps brachii (BB), and triceps brachii (TB) (Fig. 6A). EMG data were recorded at a sampling frequency of 3000Hz (TeleMyo 2400 G2, Noraxon, AZ, USA) using surface electrodes (bipolar silver/silver chloride electrodes) attached to the participant’s skin. Before starting the experiment trials, baseline EMG of muscles at rest and during maximum voluntary contraction (MVC) were recorded based on a previously described method [29]. We used a 7-camera infrared motion capture system (OptiTrack Prime 13) to track the three-dimensional positions of reflective marker clusters that were placed on the participant’s upper back, arm, forearm, and hand [30]. Marker cluster position data were recorded at 120 Hz.

#### 3) Data Processing

The *MotionMonitor* software (V8.0, Innovative Sports Training, Inc., Chicago, IL, USA) was used to compute and resample (i.e. 3000 Hz) the shoulder elevation and plane of elevation angles from the marker cluster position data. Shoulder elevation angle (sagittal and frontal trials only) or elevation plane angle (horizontal trials only) was considered for further analyses since movements were predominantly along these degrees of freedom. The EMG data for each movement repetition were divided into positive elevation/adduction, and negative elevation/abduction phases based on the kinematics. The raw EMG data was band-pass filtered at 10-500 Hz, fullwave rectified, and low pass filtered with a 4^th^ order Butterworth filter at a cut-off frequency of 3 Hz. For each participant, the processed EMG for each muscle were normalized by the maximum processed-EMG value recorded from the respective muscle during MVCs. The root mean square (nRMS) and peak (nPeak) of the normalized EMG were calculated and averaged across five middle repetitions (i.e. out of 10 repetitions) for each trial and movement phase [31, 32].

#### 4) Statistical Analysis

Paired Student’s t-tests were used to compare EMG between exoskeleton conditions (α=0.05). Since the biomechanics experiment was preliminary, we did not correct for multiple tests so that we could identify potential trends in differences between exoskeleton conditions. All statistical tests were performed using IBM SPSS software (V25, IBM Corporation, Armonk, NY, USA)

## IV. RESULTS

### A. Mechanical Performance

Under quasi-static conditions, the constant-torque spring was able to produce a nearly constant torque (0.77 ± 0.02 *Nm*) that was in the range of nominal torque (0.84 ± 0.08 *Nm*) for the stretching phase (Fig. 7). The torque generated by the spring during the unloading phase was 0.67 ± 0.02 *Nm* which was 20% lower than the nominal torque.

**Fig. 7.**
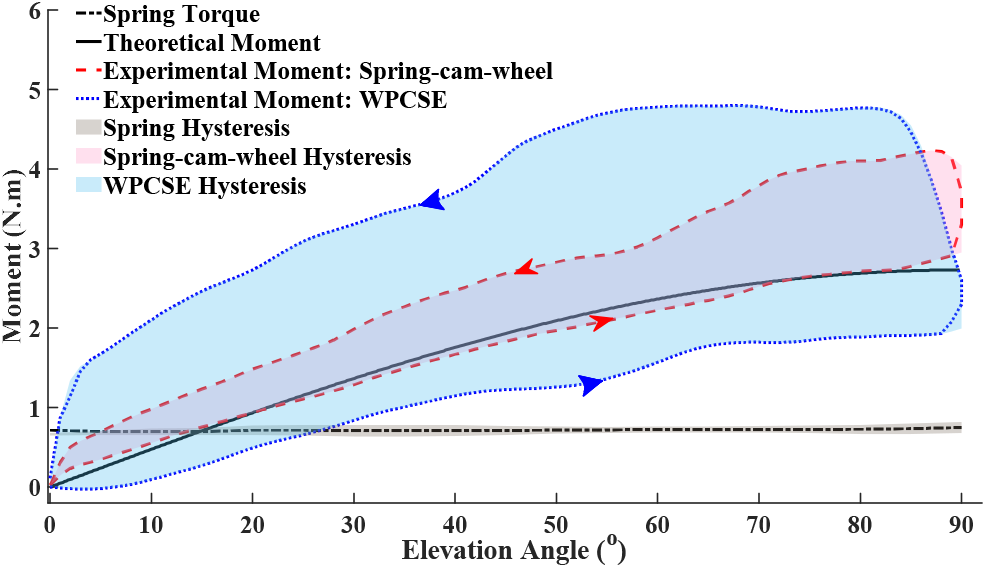
Mechanical performance of the constant-torque spring, spring-cam-wheel system, and WPCSE. The shaded regions indicate the degree of hysteresis, which is the difference in torque or moment between the positive and negative shoulder elevation phases.

For the spring-cam-wheel assembly, the measured moment matched reasonably well with the theoretical moment, especially during simulated positive shoulder elevation. This was expected because we optimized the design parameters of the cam-wheel based on the torque value that spring generates during its unloading phase. However, a larger moment, up to 50% larger than the theoretical moment, was required during simulated negative shoulder elevation to stretch the spring. The difference in moment between simulated positive and negative shoulder elevation phases was even greater for the WPCSE (Fig. 7, blue area); the measured WPCSE moment was up to 30% lower than the theoretical moment during simulated positive shoulder elevation, and up to 120% higher than the theoretical moment during simulated negative shoulder elevation.

### B. Biomechanical Performance

During the positive shoulder elevation/adduction phase, the nRMS EMG values of several muscles were lower with the WPCSE than without (Fig. 8, left column). The difference in nRMS EMG between exoskeleton conditions was most pronounced for the MD (30%), PD (25%), and ISP (27%) muscles for positive sagittal movements; AD (22%), MD (30%), ISP (27%), and TRAP (35%) muscles for positive frontal movements; and MD (9%) for horizontal adduction movements. During the negative elevation/abduction phase (Fig. 8, right column), the values of nRMS EMG were similar between exoskeleton conditions, except that the nRMS EMG value for TRAP was significantly lower (31%) with the WPCSE than without.

**Fig. 8.**
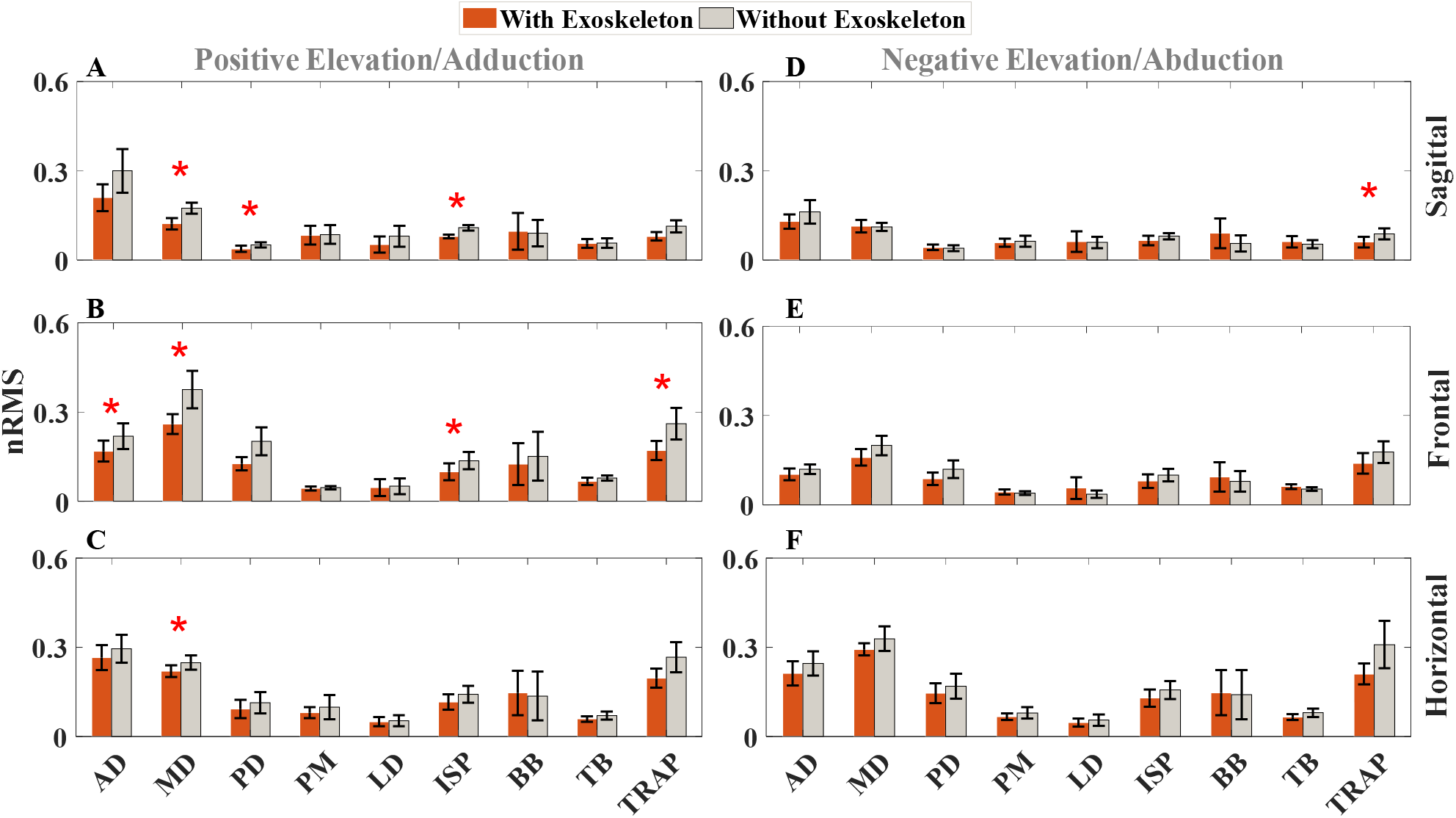
Neuromuscular activity (nRMS) of shoulder complex during shoulder elevation and horizontal abduction/adduction movements with and without WPCSE. Left column depicts the positive elevation/adduction phase in A) Sagittal plane. B) Frontal plane. C) Horizontal plane. Right column depicts the negative elevation/abduction phase in D) Sagittal plane. E) Frontal plane. F) Horizontal plane. Each bar shows the mean of nRMS across volunteers. Error bars represent the standard error. * stands for statistically significant differences.

Similarly, during the positive elevation/adduction phase, nPeak EMG values tended to be lower for trials with the WPCSE (Fig. 9). nPeak EMG was significantly lower with the WPCSE than without for the MD (23%), PD (25%), LD (33%), and ISP (21%) muscles during positive sagittal movements; for AD (14%), LD (10%), and TRAP (27%) muscles for positive frontal movements, and for the ISP (20%) muscle only during horizontal adduction movements. During negative shoulder elevation/abduction movements, nPeak EMG was similar between exoskeleton conditions for all muscles except ISP during the horizontal movement (17% lower with the WPCSE than without).

**Fig. 9.**
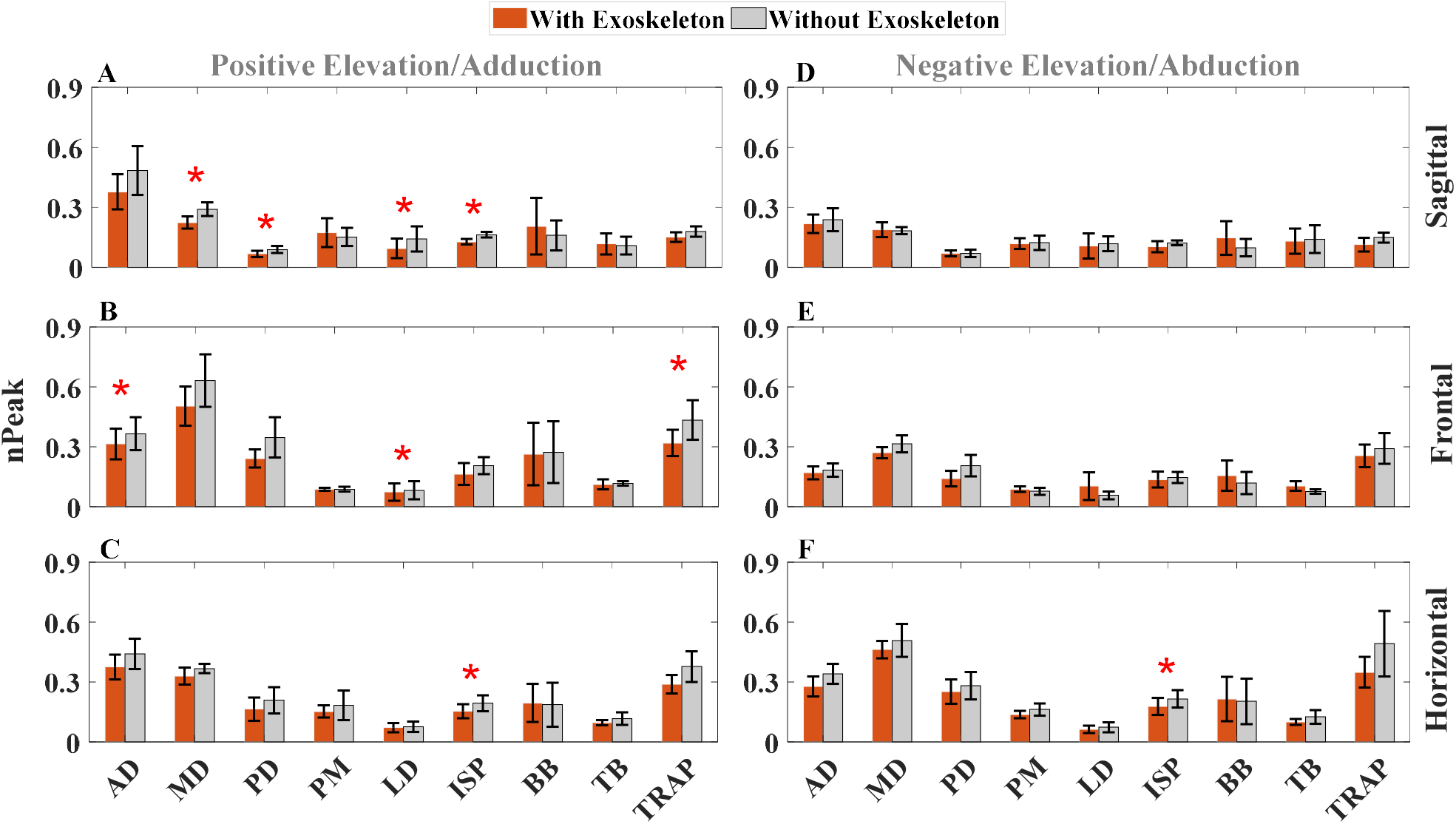
Neuromuscular activity (nPeak) of shoulder complex during shoulder elevation and horizontal abduction/adduction movements with and without WPCSE. Left column depicts the positive elevation/adduction phase in A) Sagittal plane. B) Frontal plane. C) Horizontal plane. Right column depicts the negative elevation/abduction phases in D) Sagittal plane. E) Frontal plane. F) Horizontal plane. Each bar shows the mean of nRMS across volunteers. Error bars represent the standard error. * stands for statistically significant differences.

## V. DISCUSSION

The mechanical performance results indicated the potential of the integrated spring-cam-wheel system to provide nonlinear, customizable mechanical assistance for positive shoulder elevation. The customized assistance is made possible specifically by the variable-radius cam, which acts as a gearing mechanism to modulate the force output of the spring. Cams are excellent design features for force modulation since the force can be modulated in a wide variety of ways based on the cam’s specific shape. Thus, cams have been used in many other passive mechanisms, including gravity-compensating passive shoulder exoskeletons [33]. Customizable assistance is an essential feature of exoskeletons since assistance requirements vary (1) across users as a function of their anthropometry and functional ability level and (2) across applications and tasks.

For both the spring-cam-wheel assembly and the WPCSE, the moment measured during the mechanical performance evaluation was larger during negative shoulder elevation (i.e. when the spring is stretched) than during positive shoulder elevation (i.e. when the spring recoils). We suspect that this so-called hysteresis behavior is primarily due to friction, for instance, at bearings and between the cables and components they wrap around. Inefficiency in the energy storage and return within the spring itself, inherent in all elastic springs, is another potential source of loss.

The hysteresis effect was more dramatic (i.e. more than 2x larger) with the WPCSE, which included the Bowden cable, than with the spring-cam-wheel assembly alone. Although Bowden cables are advantageous in wearable robotics applications in terms of design flexibility and remote actuation, they clearly introduce considerable friction [34]. This friction, which originates from the contact between the inner wire and outer casing of the Bowden cable, is considered to be a nonlinear function of multiple geometric and material properties [35]. In the design of the WPCSE, we tried to minimize the Bowden cable friction by (1) fixing the cable’s routing path using self-aligning spherical bearings and (2) placing the bearings so that the cable followed the most direct and shortest path from the cam-wheel to the shoulder and arm cuff. However, the Bowden cable still appeared to be the main cause for the hysteresis behavior in the WPCSE.

As demonstrated by our preliminary biomechanical evaluation, the mechanical assistance applied by the WPCSE effectively compensated for the gravity moment at the shoulder. Even when only compensating for one-fourth of the gravity moment, the WPCSE promisingly reduced the average (ranging from 9% to 35%) and peak (ranging from 10% to 33%) neuromuscular activity of several muscles crossing the shoulder during positive shoulder elevation/adduction. Our results are consistent with several previous studies that also showed that varying levels of gravity compensation at the shoulder can reduce muscle activity [8, 9, 36, 37].

Though the WPSCE reduced activity of several muscles, reductions were most significant for the deltoid and rotator cuff muscles. The deltoid mainly contributes to positive shoulder elevation and, thus, plays a major role in compensating for gravity at the shoulder [38]. Infraspinatus, a superficial rotator cuff muscle that contributes to stabilization of the glenohumeral joint [38], also had lower activity with the WPCSE than without. Though we did not measure their activity directly, the supraspinatus and teres minor rotator cuff muscles likely had lower activity with the WPCSE, too. This can be inferred from the trapezius muscle whose activity, which was lower with the WPCSE, was previously shown to be positively correlated with the activity of the supraspinatus and teres minor muscles [39]. In future studies, we plan to measure neuromuscular activity in all rotator cuff muscles, given their importance in shoulder motor function and stability.

The level of mechanical assistance generated by the WPCSE is a key design parameter that is expected to have a strong influence on neuromuscular activity, joint loads, and motor function at the shoulder. A desirable level of assistance for the WPCSE would be one that maximally assists and minimally resists user-generated moments. For example, though a high level of gravity compensation at the shoulder would maximally assist positive shoulder elevation movements, it might also require the user to generate higher negative shoulder elevation moments to overcome the gravity compensation and lower the arm. More research is needed to investigate the relationship between exoskeleton assistance level and biomechanical performance, which will inform the future design and prescription of wearable passive shoulder exoskeletons.

Interestingly, the WPCSE did not substantially affect muscle activity during negative shoulder elevation. This indicates that the participants were likely able to lower their arm using mostly gravity, even though the moment generated by the WPCSE during negative shoulder elevation was considerably higher than we designed it to be. The WPCSE even reduced the activity of the trapezius and infraspinatus muscles. This result, promisingly, contrasts our previous computational study, which predicted higher activity in several muscles during negative shoulder elevation [27]. One possible explanation is that, during such movements, the WPCSE offloads muscles that contribute to positive shoulder elevation that are eccentrically contracted to control the shoulder’s angular velocity. Additionally, participants may adapt their kinematics in subtle ways to avoid increasing muscle activity. We will test these hypotheses in a future, more comprehensive biomechanical evaluation of the WPCSE.

## VI. Future Work & Conclusion

To successfully translate assistive technology to people with disability, it is critical to understand its effectiveness, usability, and biomechanical interaction with humans. As a first step toward accomplishing this goal, we quantitatively evaluated the mechanical and biomechanical performance of our WPCSE prototype. Our results showed that the WPCSE, compensating for a modest one-fourth of the gravity moment at the shoulder, reduced neuromuscular activity of several muscles crossing the shoulder without increasing activity of any muscles during either positive or negative shoulder elevation and horizontal adduction/abduction movements. However, our mechanical evaluation revealed aspects of the design that limit the assistance the WPCSE could provide. In our future work, we will test different WPCSE assistance levels and identify a range of assistance that most enhances shoulder motor function and biomechanics, without impeding movements. More comprehensive biomechanical studies will be performed to assess the WPCSE for more biomechanical parameters (e.g. joint kinematics), more participants (both able-bodied subjects and people with shoulder disability), and more movements that typify activities of daily living. Finally, several design refinements need to be made, especially those that reduce friction in the system.

## Acknowledgement

This study was supported by start-up funds from the Department of Mechanical, Aerospace and Biomedical Engineering at the University of Tennessee. The authors thank Vulcan Spring & Manufacturing for providing the constanttorque springs.

